# Serotonergic Psychedelics LSD & Psilocybin Increase the Fractal Dimension of Cortical Brain Activity in Spatial and Temporal Domains

**DOI:** 10.1101/517847

**Authors:** TF Varley, R Carhart-Harris, Leor Roseman, David K Menon, EA Stamatakis

**Affiliations:** Division of Anaesthesia, School of Clinical Medicine, University of Cambridge, UK; Department of Clinical Neurosciences, School of Clinical Medicine, University of Cambridge, UK; Centre for Neuropsychopharmacology, Department of Medicine, Imperial College London, London, UK; Computational, Cognitive and Clinical Neuroscience Laboratory, Department of Medicine, Imperial College London, London, UK; Department of Psychological & Brain Sciences, Indiana University, Bloomington, IN, USA

**Keywords:** Complexity, Consciousness, Criticality, Entropy, fMRI, Fractal, LSD, Networks, Psilocybin, Psychedelic

## Abstract

Psychedelic drugs, such as psilocybin and LSD, represent unique tools for researchers investigating the neural origins of consciousness. Currently, the most compelling theories of how psychedelics exert their effects is by increasing the complexity of brain activity and moving the system towards a critical point between order and disorder, creating more dynamic and complex patterns of neural activity. While the concept of criticality is of central importance to this theory, few of the published studies on psychedelics investigate it directly, testing instead related measures such as algorithmic complexity or Shannon entropy. We propose using the fractal dimension of functional activity in the brain as a measure of complexity since findings from physics suggest that as a system organizes towards criticality, it tends to take on a fractal structure. We tested two different measures of fractal dimension, one spatial and one temporal, using fMRI data from volunteers under the influence of both LSD and psilocybin. The first was the fractal dimension of cortical functional connectivity networks and the second was the fractal dimension of BOLD time-series. We were able to show that both psychedelic drugs significantly increased the fractal dimension of functional connectivity networks, and that LSD significantly increased the fractal dimension of BOLD signals, with psilocybin showing a non-significant trend in the same direction. With both LSD and psilocybin, we were able to localize changes in the fractal dimension of BOLD signals to brain areas assigned to the dorsal-attentional network. These results show that psychedelic drugs increase the fractal character of activity in the brain and we see this as an indicator that the changes in consciousness triggered by psychedelics are associated with evolution towards a critical zone.

**Author Summary:** The unique state of consciousness produced by psychedelic drugs like LSD and psilocybin (the active component in magic mushrooms) are potentially useful tools for discovering how specific changes in the brain are related to differences in perception and thought patterns. Past research into the neuroscience of psychedelics has led to the proposal of a general theory of brain function and consciousness: the Entropic Brain Hypothesis proposes that consciousness emerges when the brain is sitting near a critical tipping point between order and chaos and that the mind-expanding elements of the psychedelic experience are caused by the brain moving closer to that critical transition point. Physicists have discovered that near this critical point, many different kinds of systems, from magnets to ecosystems, take on a distinct, fractal structure. Here, we used two measures of fractal-quality of brain activity, as seen in fMRI, to test whether the activity of the brain on psychedelics is more fractal than normal. We found evidence that this is the case and interpret that as supporting the theory that, psychedelic drugs are move the brain towards a more critical state.

## 1 Introduction

Since the turn of the century, there has been a renewal of interest in the science of serotonergic psychedelic drugs (LSD, psilocybin, mescaline, etc.), both in terms of possible medical applications of these drugs [1, 2], and what they might tell us about the relationship between activity in the brain and the phenomenological perception of consciousness [3, 4]. For those interested in the relationship between activity in the brain and consciousness, psychedelic drugs are particularly useful, as volunteers under the influence of a psychedelic are still able to report the nature of their experience and recall it even after returning to normal consciousness. This contrasts favourably with the other class of drugs commonly used to explore consciousness: anaesthetics, which by the very nature of their effects, make it difficult to gather first-person experiential data from a volunteer [5]. The subjective experience of the psychedelic state is associated with radical alterations to both internal and external senses, including visual distortions, vivid, complex closed-eye imagery, alterations to the sense of self, emotional extremes of euphoria and anxiety, and in extreme cases, psychosis-like effects [6]. The psychedelic experience can also have profound personal, and even spiritual or religious character [7, 8], which has made them central to the religious practices of many cultures around the world [9]. In this way, the study of the psychedelic state can inform not just the question of why consciousness emerges, but also the origins of some of the most quintessentially human psychological experiences.

Neuroimaging studies using fMRI and MEG have suggested that the experiential qualities of the psychedelic state can be explained, in part, by the effects these drugs have on the entropy of brain activity: a theory known as the Entropic Brain Hypothesis (EBH) [10, 4]. The EBH posits that during normal waking consciousness, activity in the brain is near, but slightly below, a critical zone between order and disorder, and that under the influence of psychedelic drugs the entropy of brain activity increases, bringing the system closer to the zone of criticality. In this context, ‘criticality’ can be thought of as similar to a phase-transition between two qualitatively different states: the sub-critical state, which is comparatively inflexible, highly ordered and displays low entropy, while the super-critical state may be highly entropic, flexible, and disorganized (this recalls a canonical model of critical processes, the Ising Model, where the critical temperature divides distinct phases, one where the magnetic spins are all aligned, and another where the spins are distributed chaotically, for review see [11]). The EBH is related to a larger theory of consciousness, known as Integrated Information Theory (IIT), which posits that consciousness is an emergent property of the integration of information in the brain [12, 13, 14] and that this mathematical formalism is categorically isomorphic to consciousness itself [15].

While it is currently impossible to directly measure the entropy of all of the activity in the whole of the brain, or the amount of information integration, there is much interest in using mathematical analysis of neuroimaging data to estimate the complexity of activity in the brain and relate that to consciousness. Studies with psilocybin have found that the patterns of functional connectivity in the brain undergo dramatic reorganization, characterized primarily by the rapid emergence and dissolution of unstable communities of interacting brain regions that do not occur in normal waking consciousness [16]. Similarly, under psilocybin, the repertoire of possible states functional connectivity networks can occupy is increased, which is interpreted as an increase in the entropy of the entire system [17]. Work on other psychedelics with pharmacology related to psilocybin has found similar results: under the influence of Ayahuasca, a psychedelic brew indigenous to the Amazon, the Shannon entropy of the degree distribution of functional connectivity networks is increased relative to normal consciousness [18] (encouragingly, the opposite effect has been shown under the conditions of sedation with propofol [19]). Analysis of MEG data from volunteers under the influence of lysergic acid diethylamide (LSD) has been shown an increase in the Lempel-Ziv complexity of the signals, which is thought to reflect increased complexity of activity in the brain [20]. LSD has also been recently shown to alter the connectome harmonics of brain networks, in a manner that suggests an increase in the complexity of network harmonics describing brain activity [21]. For a comprehensive review of the current state of psychedelic research into the EBH see *The Entropic Brain - Revisited* [4].

While a core element of the EBH is the theory that the psychedelic experience moves the brain closer to the zone of criticality, many of the measures that have been tested so far do not address the phenomena of criticality directly. These measures usually test where the brain falls on a unidimensional axis of order vs. randomness. Lempel-Ziv complexity [20], nodal entropy [18, 19] and the entropy of possible states [17], all describe a movement towards increased randomness and disorder, which is consistent with the entropic predictions of the EBH, but not necessarily informative about the relative proximity to the zone of criticality. In these analyses, a completely random system would score maximally high on complexity (for instance a completely random time-series would have a normalized Lempel-Ziv score of unity, which is the upper bound of the measure) however it is nearly impossible to imagine how a living brain could output totally random data, and such a brain would most likely not be conscious. While these analyses are interesting and have clearly been fruitful, they paint a limited picture of the brain as a complex system, and don’t directly test the central thesis of the EBH. To date, the only study that has directly addressed the criticality aspect of the EBH is the study of LSD and connectome harmonics [21], although other studies have found evidence of scale-free, power-law behaviour generally thought to be indicative of critical phenomena [22]. To address the relative lack of studies testing criticality directly, in this paper, we propose the fractal dimension of brain activity as a novel measure of complexity that provides insights into the criticality of the psychedelic state, as well as providing a measure of ‘complexity’ that is related to, but distinct from, the entropic measures described above.

Fractals are ubiquitous in nature and dramatic visualizations of colourful constructs like the Mandelbrot set have even permeated popular culture [23]. Psychedelic culture in particular shows a strong affinity for fractal patterns, as much of the imagery experienced under the influence of psychedelics is described as fractal in character. Fractals are defined by the property of having a non-integer dimension, which can be naively thought of as how ‘rough’ or ‘complex’ the shape in question is, or slightly more formally, the extent to which it maintains symmetry across different scales [24]. This is commonly known as ‘self-similarity,’ and can be intuitively understood as the invariance of appearance across scales: for example, the pattern of small creeks flowing together can resemble the pattern of large rivers carrying orders of magnitude more water [25]. In systems that display self-organizing criticality, as the system naturally evolves towards a critical point, its spatial structure will tend to take on increasingly fractal character that can be described in terms of fractal dimension [26, 27, 28], and in systems which can be ‘tuned’ to a critical state (such as the Ising model, which has been explored as a model of critical brain activity [29, 30, 31]), fractal structures emerge near the critical point [32]. If, under the influence of a psychedelic, the brain is moving closer towards a state of criticality, as the EBH posits, then we might expect any fractal character in brain activity to become more pronounced. There is some evidence of a symmetrical effect when consciousness is lost: in states of sleep and drug-induced anaesthesia, the fractal dimension of brain activity drops significantly, with the exception of REM sleep, during which the fractal dimension rises again [33, 34]. As REM sleep is the state of sleep when the greatest quantity of phenomonological experience takes place (in the form of dreams), this suggests that the fractal dimension of brain activity is related to the ‘quantity’ of experiential consciousness available to an individual. Similarly, in rats, during ketamine-induced anaesthesia the fractal dimension of brain activity is significantly higher in key-brain regions associated with consciousness when compared with anaesthesia induced by other drugs [35], and as ketamine is known to induce vivid, dream-like states of consciousness at high doses [36], which comports with the REM sleep finding.

There has been considerable interest in applying techniques of fractal analysis to questions in neuroscience and considerable evidence has mounted that both the physical structure of the brain itself, and the patterns of activity measured by neuroimaging paradigms display pronounced fractal character [37, 38]. Changes to the fractal dimension of brain structures are associated with changes in cognition and clinically significant diagnosis, such as schizophrenia and obsessive-compulsive disorder [39], intelligence [40], Alzheimer’s disease [41], and ageing [42]. There is some preliminary evidence that cortical functional connectivity networks display fractal character, both during rest and tasks [43] and that this fractal character plays an important role in regulating how information is propagated through the brain [44].

While fractal dimension is usually though to encode complexity in terms of self-similarity rather than entropy directly, there is a connection between the two values: fractal dimension is related to Renyi entropy, which is itself a generalization of the classical measure of Shannon entropy [45]. Computational models have shown that as the fractal dimension of a shape rises, so does the associated Renyi entropy [46]. Another measure, the information dimension, relates the fractal dimension to the information content of a fractal at different scales [47, 48]. Based on these findings, and the results reported by Bak et al., (1987), we propose that the fractal dimension is a natural metric by which to test the EBH, for several reasons. First, unlike other metrics of entropy, fractals are intimately related to the phenomena of criticality, which is predicted to be significant for consciousness, and the fractal dimension encodes information relevant to a system’s evolution towards criticality. Second, in this context, they are a novel method of describing the behaviour of the brain as a complex system and so give information beyond the axis of order versus randomness. Finally, despite the differences between the measure of fractal dimension and classical entropy, the two are related in some fundamental ways. The fractal dimension sits at a sweet spot of not being so radical that it cannot be related to previous results, while still being novel enough to open the door to new and informative avenues of study.

To test the relationship between the fractal dimension of activity of brain and consciousness, we used fMRI data from subjects under the influence of either LSD or psilocybin, provided by the Psychedelic Research Group at Imperial College London. From this data, we created 1000-node functional connectivity networks and performed a network-specific variation of the box-counting algorithm [49] to extract the fractal dimension. We also used a second measure, the Higuchi fractal dimension [50], to test the temporal fractal dimension of BOLD time-series. These two measures capture two axes on which the complexity of brain activity might be measured: spacial (network fractal dimension) and temporal (Higuchi fractal dimension). If the psychedelic state is associated with a movement towards a critical zone associated with increased fractal character, we would expect to see this when examined on multiple measures, and so these two measures serve as internal validation for each-other. While the network fractal dimension is not spacial in the way, for example, a 2-dimensional box-counting analysis of activity at the cortical surface would be, it does return insight into how information processing may be distributed across multiple, spatially distinct brain regions.

## 2 Materials & Methods

### 2.1 Ethics Statement

The data analyzed here have been reported in previous studies [59, 58]. Both studies described herein were approved by a UK National Health Service research ethics committee, and the researchers complied with all relevant regulations and ethical guidelines, including data privacy and participant informed consent.

### 2.2 Calculating Network Fractal Dimension

When calculating the fractal dimension of a naturally occurring system, researchers commonly use a box-counting algorithm, which is an accessible and computationally tractable method that captures the distribution of elements across multiple scales [24]. Intuitively, the box-counting dimension defines the relationship between a measured quality of a shape in space, and the metric used to measure it. The canonical example is the question of how long the coastline of Britain is [51]. If one wishes to measure the length of Britain’s coast, they could estimate it by calculating the number of square boxes *N*_*B*_(*l*_*B*_), of a given width *l*_*B*_, that are necessary to tile the entire coastline. For very large values of *l*_*B*_, *N*_*B*_(*l*_*B*_) will be small, while as the value of *l*_*B*_ decreases, *N*_*B*_(*l*_*B*_) will asymptotically approach some value. If the shape being tiled is a fractal, then:

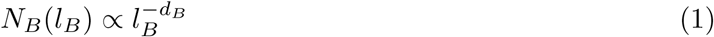

Where *d*_*B*_ is the box-counting dimension. Algebraic manipulation shows that *d*_*B*_ can be extracted by linear regression in log-log space as:

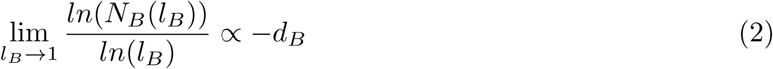

A similar logic is used when calculating the box-counting dimension of a graph. For a graph *G* = (*V, E*), a box with diameter *l*_*B*_ defines a set of nodes *B* ⊂ *V* where for every pair of nodes *v*_*i*_ and *v*_*j*_ the distance between them *l*_*ij*_ < *l*_*B*_. Here, the distance between two nodes *v*_*i*_, *v*_*j*_ is the graph geodesic between the vertices: the number of edges in the shortest path between them. To quantify the fractal dimension of the functional connectivity networks, a box counting method, the Compact Box Burning algorithm (CBB) [49], was used to find *N*_*B*_(*l*_*B*_) for a range of integer *l*_*B*_ values 1..10. If *G* has fractal character, a plot of *ln*(*N*_*B*_(*l*_*B*_)) vs. *ln*(*l*_*B*_) should be roughly linear, with a slope of *-d*_*B*_. Unfortunately, because of the logarithmic relationship between box-size and fractal dimension, exponentially higher resolutions are required to achieve modest increases in the accuracy of the measured fractal dimension. Computational explorations, where a box-counting method is used to approximate a fractal dimension that has already been solved analytically, show that the box-counting dimension converges to the true dimension with excruciating slowness [52], necessitating the highest-resolution parcellation that is computationally tractable.

It should be noted that there has been much discussion surrounding the appropriateness of this method for describing the presence (or absence) of power-laws in empirical data [53]. We chose the above-described method for a few reasons: the first was to remain as consistent as possible with the method used in previous analysis of the fractal dimension of human FC networks [44, 43], the second was because of the tractability of the analysis, and finally, the relatively small size of the network forced a limited range of box sizes *l*_*B*_ (approximately a single order of magnitude), which precluded the use of larger, more data-driven analyses. We stress that, given the ongoing discussion around the optimal way to find power-law relationships, the results reported here should not be interpreted as an unambiguous claim of incontrovertible proof that such a power-law relationship holds here - rather a preliminary result to establish the possibility that fractal topologies and brain dynamics may be related to the maintenance of consciousness.

The implementation of the CBB was provided as open-source code by the Mackse lab, and can be found at: http://www-levich.engr.ccny.cuny.edu/webpage/hmakse/software-and-data/

### 2.3 Calculating BOLD Time-Series Fractal Dimension

To calculate the temporal fractal dimension, we used the Higuchi method for calculating the self-similarity of a one-dimensional time-series, an algorithm widely used in EEG and MEG analysis [54]. The original method is recorded in detail in the original paper [50], but will be briefly described here. The algorithm takes in a time-series *X*(*t*) with *N* individual samples, defined as:

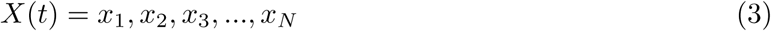

In this case, every *X*(*t*) corresponds to one Hilbert-transformed BOLD time-series *H*(*t*) extracted from our functional brain scans (details below). Hilbert-transforming was chosen to be consistent with previously-reported studies of time-series complexity and consciousness [55, 56, 20]. From each time-series *X*(*t*), we create a new time-series 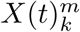, defined as follows:

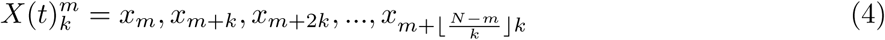

where *m* = 1, 2, *…, k*.

For each time-series 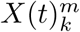 in *k*_1_, *k*_2_, *…k*_*max*_, the length of that series, *L*_*m*_(*k*), is given by:

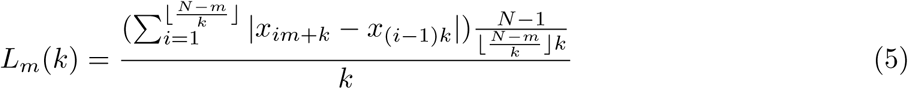

We then define the average length of the series ⟨ *L*(*k*) ⟩, on the interval [*k, L*_*m*_(*k*)] as:

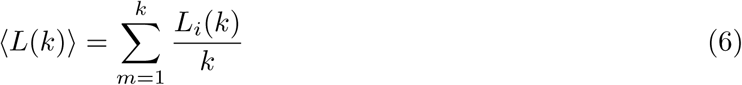

If our initial time-series *X*(*t*) has fractal character, then ⟨ *L*(*k*) ⟩ ∝ *k*^*-D*^. As with the procedure for calculating the network fractal dimension, the algorithm iterates through values of *k* from 1*…k*_*max*_ and calculates *ln*(⟨ *L*(*k*) ⟩) vs. *ln*(*k*^*-*1^), extracting *D* by linear regression. The various values of *k* can be thought of as analogous to the various values of *l*_*B*_ used to calculate the network fractal dimension. The Higuchi algorithm requires a pre-defined *k*_*max*_ value as an input, along with the target time-series. This value is usually determined by sampling the results returned by different values of *k*_*max*_ and selecting a value based on the range of *k*_*max*_ where the fractal dimension is stable. For the psilocybin and LSD datasets, we sampled over a range of powers of two (2, *…,* 128). Due to the comparably small size of BOLD time-series (100 entries for the psilocybin dataset and 434 entries for the LSD dataset), the range of *k*_*max*_ values that our algorithm could process without returning an error was limited. We ultimately decided on *k*_*max*_ = 64 for the LSD dataset and *k*_*max*_ = 32 for the psilocybin dataset.

The implementation we used was from the PyEEG toolbox [57], downloaded from the Anaconda repository.

### 2.4 Data Acquisition & Preprocessing

Both the LSD data and the psilocybin data were provided by the Psychedelic Research Group at Imperial College London, having already been preprocessed according to their specifications.

#### 2.4.1 LSD Data

The data acquisition protocols and preprocessing pipelines were described in detail in a previous paper [58], so we will describe them in brief here. 20 healthy volunteers underwent two scans, 14 days apart. On one day they were given a placebo (10-mL saline) and on the other they were given an active dose of LSD (75 *µ*g of LSD in 10-mL saline). BOLD scanning consisted of three seven minute eyes closed resting state scans. The first and third scans were eyes-closed, resting state without any in-ear auditory stimulation (music), and these were what were used in this report.

Anatomical imaging was performed on a 3T GE HDx system. These were 3D fast spoiled gradient echo scans in an axial orientation, with field of view = 256 × 256 × 192 and matrix = 256× 256× 129 to yield 1mm isotropic voxel resolution. TR/TE = 7.9/3.0ms; inversion time = 450ms; flip angle = 20^°^ BOLD-weighted fMRI data were acquired using a gradient echo planer imaging sequence, TR/TE = 2000/35ms, FoV = 220mm, 64×64 acquisition matrix, parallel acceleration factor = 2, 90^°^ flip angle. Thirty five oblique axial slices were acquired in an interleaved fashion, each 3.4mm thick with zero slice gap (3.4mm isotropic voxels). The precise length of each of the two BOLD scans was 7:20 minutes. One subject aborted the experiment due to anxiety and four others were excluded for excessive motion (measured in terms of frame-wise displacement).

The following pre-processing stages were performed: removal of the first three volumes, de-spiking (3dDespike, AFNI), slice time correction (3dTshift, AFNI), motion correction (3dvolreg, AFNI) by registering each volume to the volume most similar to all others, brain extraction (BET, FSL); 6) rigid body registration to anatomical scans, non-linear registration to 2mm MNI brain (Symmetric Normalization (SyN), ANTS), scrubbing (FD = 0.4), spatial smoothing (FWHM) of 6mm, band-pass filtering between [0.01 to 0.08] Hz, linear and quadratic de-trending (3dDetrend, AFNI), regressing out 9 nuisance regressors (all regressors were bandpass-filtered using the same range described above).

#### 2.4.2 Psilocybin Data

The data acquisition protocols and preprocessing pipelines were described in detail in a previous paper [59], so we will describe them in brief here. Fifteen healthy volunteers were scanned. Anatomical and task-free resting state scans (each lasting 18 minutes) were taken. Solutions were infused manually over 60 s, beginning 6 min after the start of each functional scan. Subjects psilocybin (2 mg in 10-mL saline) in the active scan. In this study we used only the psilocybin-positive scan, comparing the pre-infusion condition to the post-infusion condition for control.

All imaging was performed on a 3T GE HDx system. For every functional scan, we obtained an initial 3D FSPGR scan in an axial orientation, with FoV = 256 × 256 × 192 and matrix = 256 × 256 × 192 to yield 1-mm isotropic voxel resolution (TR/TE = 7.9/3.0 ms; inversion time = 450 ms; flip angle = 20^°^). BOLD-weighted fMRI data were acquired using a gradient-echo EPI sequence, TR/TE 3000/35 ms, field-of-view = 192 mm, 64 × 64 acquisition matrix, parallel acceleration factor = 2, 90^°^ flip angle. Fifty-three oblique-axial slices were acquired in an interleaved fashion, each 3 mm thick with zero slice gap (3 × 3 × 3-mm voxels). A total of 240 volumes were acquired.

All data was preprocessed using the following pipeline: de-spiking, slice time correction, motion correction to best volume, brain extraction using the BET module in FSL, registration to anatomy (using FSL BBR), registration to 2mm MNI (ANTS), scrubbing (FD=0.4), smoothing with a 6mm kernel, bandpass filtering [0.01-0.08 Hz], linear and quadratic detrending, regression of 6 motion regressors and 3 nuisance regressors (all of the regressors were not smoothed and were bandpassed with the same filters). At the suggestion of the original research team that provided the data, six volunteers were excluded from the analysis for excessive motion.

### 2.5 Formation of Functional Connectivity Networks

BOLD time-series data were extracted from each brain in CONN (CONN is a collection of SPM/MATLAB scripts with a GUI designed for easy manipulation of fMRI, MEG, and EEG data. It is available at http://www.nitrc.org/projects/conn) [60] and the cerebral cortex was segmented into 1000 distinct ROIs, using the “Schaefer Local/Global 1000 Parcellation” [61] (https://github.com/ThomasYeoLab/CBIG/blob/maste/Schaefer2018LocalGlobal/Parcellations/MNI/Schaefer2018_1000Parcels_7Networks_order_FSLMNI152_1mm.nii.gz) Due to the slow-convergence of Eq. 2, and the necessity of having a network with a wide enough diameter to accommodate a sufficiently wide range of box-sizes (if *l*_*B*_ is greater than or equal to the diameter of the network, then *N* (*l*_*B*_) is trivially one), we attempted to strike an optimal balance between network resolution and computational tractability.

Every time-series *F* (*t*) was first transformed by taking the norm of the Hilbert transform of each time-series, to ensure an analytic signal and keep the signals consistent with the Higuchi fractal dimension analysis.

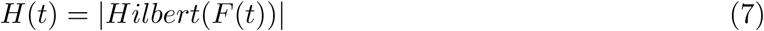

Pearson Correlation was chosen largely due to it’s wide use in the field and ease of interpretation. While more exotic, nonlinear similarity functions exist (normalized mutual information, information-based similarity, etc), for a prospective study of this sort, use of a well-characterized, linear function was appropriate, although future studies might explore the effect of different functions on large network topology. The resulting time-series *H*(*t*) was then correlated against every other time-series, using the Pearson Correlation, forming a matrix *M* such that:

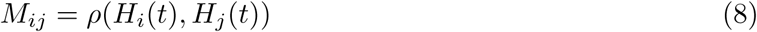

No significance testing was done (every *ρ* was included, regardless of whether it met some arbitrary *α* value or not), because significance filtering would result in an uneven distribution of edges and degrees between graphs that may have effected the analysis. Due to the high thresholding, the vast majority of weak, or potentially spurious connections were likely removed anyway. The correlation matrix has a series of ones that run down the diagonal, corresponding the correlation between each timeseries and itself which, if treated directly as a graph adjacency matrix, would produce a graph where each node had exactly one self-loop in addition to all it’s other connections. To correct for this, the matrices were filtered to remove self-loops by turning the diagonal of ones to zeros, ensuring simple graphs:

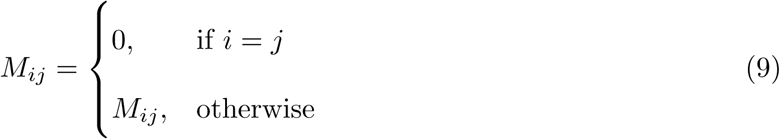

Finally, the matrices were binarized with a 95% threshold, such that:

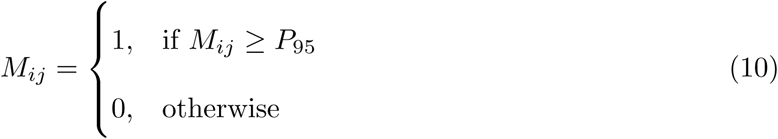

The thresholding procedure was passed over all entries in the matrix, regardless of whether they were positive or negative, and any surviving edges became ones. The practical effect of such stringent thresholding is that only positive values survived, and including the negative values drove down the minimum edge weight that survived thresholding, resulting in a marginally less sparse network than what might have occurred if negative values had been thrown out prior to thresholding. While binarization does throw out information, the CBB algorithm that we used does not factor edge weight into whether two nodes constitute members of the same box. A 95% threshold was chosen based on the findings of Gallos et al., who showed that functional connectivity networks only display fractal character at high thresholds (see Introduction). All surviving values *M*_*ij*_ < 0 ⟼ 0. The results could then be treated as adjacency matrices defining functional connectivity graphs, where each row *M*_*i*_ and column *M*_*j*_ corresponds to an ROI in the initial cortical parcellation, and the connectivity between all nodes is given by Eq. 3. To see samples of the binarized adjacency matrices, and the associated graphs see Figure 1.

**Figure 1:**
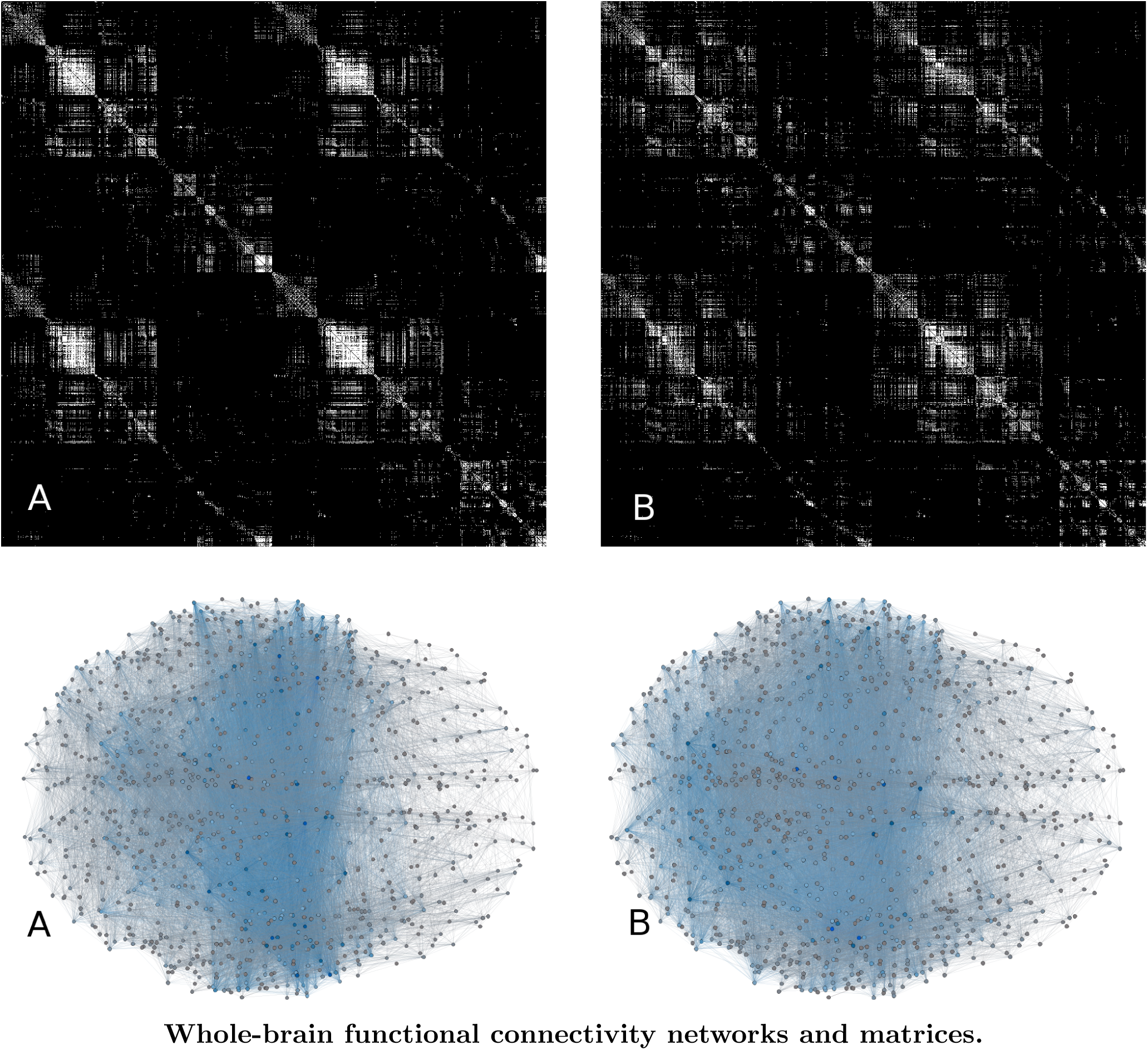
Two binarized, 1000-ROI adjacency matrices from a single subject, and their associated functional connectivity graphs (A ⟼ A, etc). In the adjacency matrices, every pixel represents an edge between two nodes: if the pixel is white, the edge exists, if black, the edge does not exist. A is the functional connectivity matrix from the placebo condition, B is the matrix from the LSD condition. While the differences in fractal character are not intuitively obvious upon visual inspection, subtle differences in the distribution of connections can be seen. When the corresponding networks are constructed, differences in gross-scale connectivity can be seen, although, as with the matrices, a change in fractal structure is not intuitively obvious. The networks are constructed using axial projections of the 3-dimensional atlas: each node is roughly at the centroid of it’s associated ROI.

#### 2.5.1 Specific-Network Analysis

To localize changes in the complexity of brain activity, individual ROIs were grouped into networks, using the mapping proposed by Yeo et al., [62]. We used the 1000 ROI parcellation with seven networks: default mode network, somato-motor network, visual network, dorsal-attentional network, ventral-attentional network, limbic network, and fronto-parietal control network. For visualization of the assignment of nodes to these networks see Figure 2. We then used the Higuchi fractal dimension method described above on each subset of regions to get a measure of the average time-series fractal dimension of each network.

**Figure 2:**
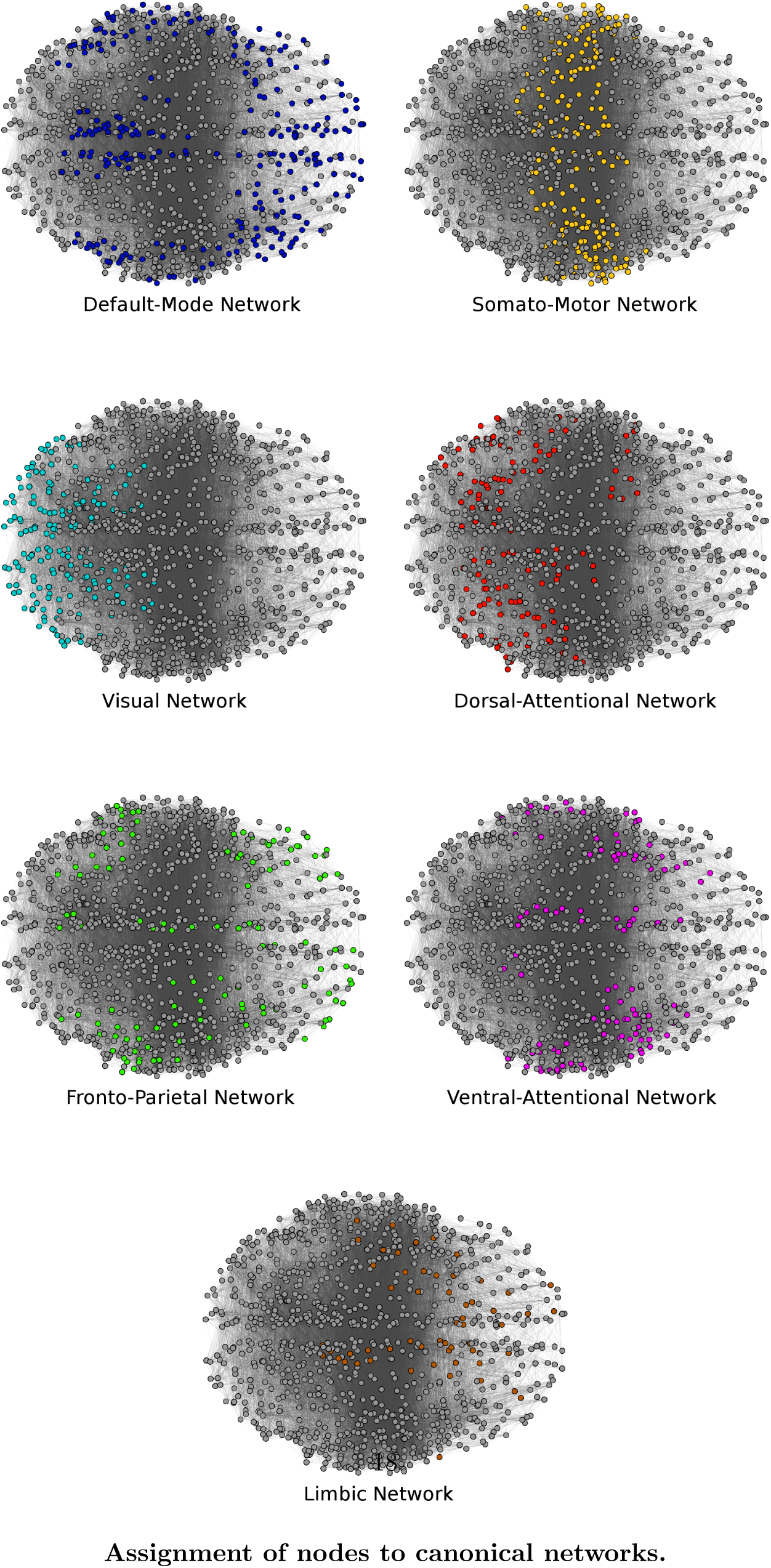
A visualization of how the 1000-node functional connectivity networks were parcellated into seven different brain regions, following the mapping described by Yeo et al., [62, 61] The specific map file is available from GitHub at https://github.com/ThomasYeoLab/CBIG/blob/master/stable_projects/brain_parcellation/Schaefer2018_LocalGlobal/Parcellations/MNI/Schaefer2018_1000Parcels_7Networks_order.txt

#### 2.5.2 Statistical Analysis

All analysis was carried out using the Python 3.6 programming language in the Spyder IDE (https://github.com/spyder-ide/spyder), using the packages provided by the Anaconda distribution (https://www.anaconda.com/download). All packages were in the most up-to-date version, with the exception of NetworkX: due to compatibility issues with the CBB code, NetworkX v. 0.36 was used. Packages used include NumPy [63], SciPy [64], and NetworkX [65]. NetworkX was used for the implementation of the CBB algorithm, NumPy was used for manipulation of adjacency matrices and arrays, SciPy was used for statistical analysis, primarily using the SciPy.Stats module. Unless otherwise specified, all the significance tests are non-parametric: given the small sample sizes and heterogeneous populations, normal distributions were not assumed. Wilcoxon Signed Rank test was used to compare drug conditions against their respective control conditions.

## 3 Results

### 3.1 LSD & Psilocybin Network Fractal Dimension

The Wilcoxon signed-rank test found significant differences, when corrected with the Benjamini-Hochberg procedure with an FDR of 5% [66], between LSD and placebo conditions (H(4), p-value = 0.001), and between the pre-infusion and post-infusion psilocybin conditions (H(6), p-value = 0.05). The mean fractal dimensions for the LSD condition was 3.37 ± 0.15, and for the associated placebo condition it was 2.939 ± 0.29. For psilocybin the mean fractal dimension was 3.52 ± 0.049, and for control it was 3.277 ± 0.372. For a plot of the relative fractal dimensions, see Figure 3. For a visualization for how the fractal dimension was calculated by linear regression for LSD see 4A and for Psilocybin, see Figure 4B.

**Figure 3:**
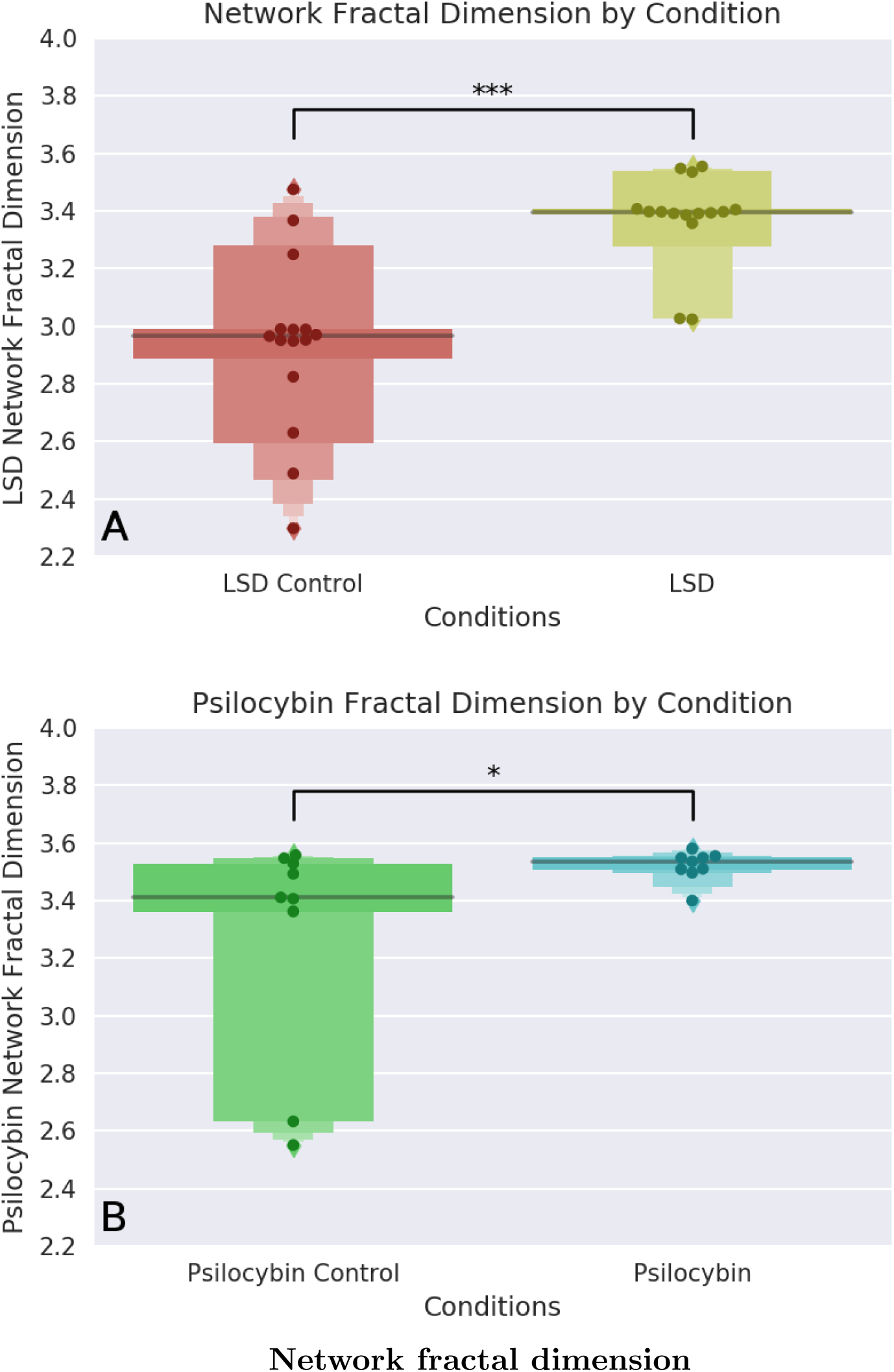
Letter-value plots of the network fractal dimensions for the two psychedelic drugs tested. Note that both psychedelic conditions show less variability compared to their respective controls. * p ≤ 0.05, ** p ≤ 0.01, *** p ≤ 0.001

**Figure 4:**
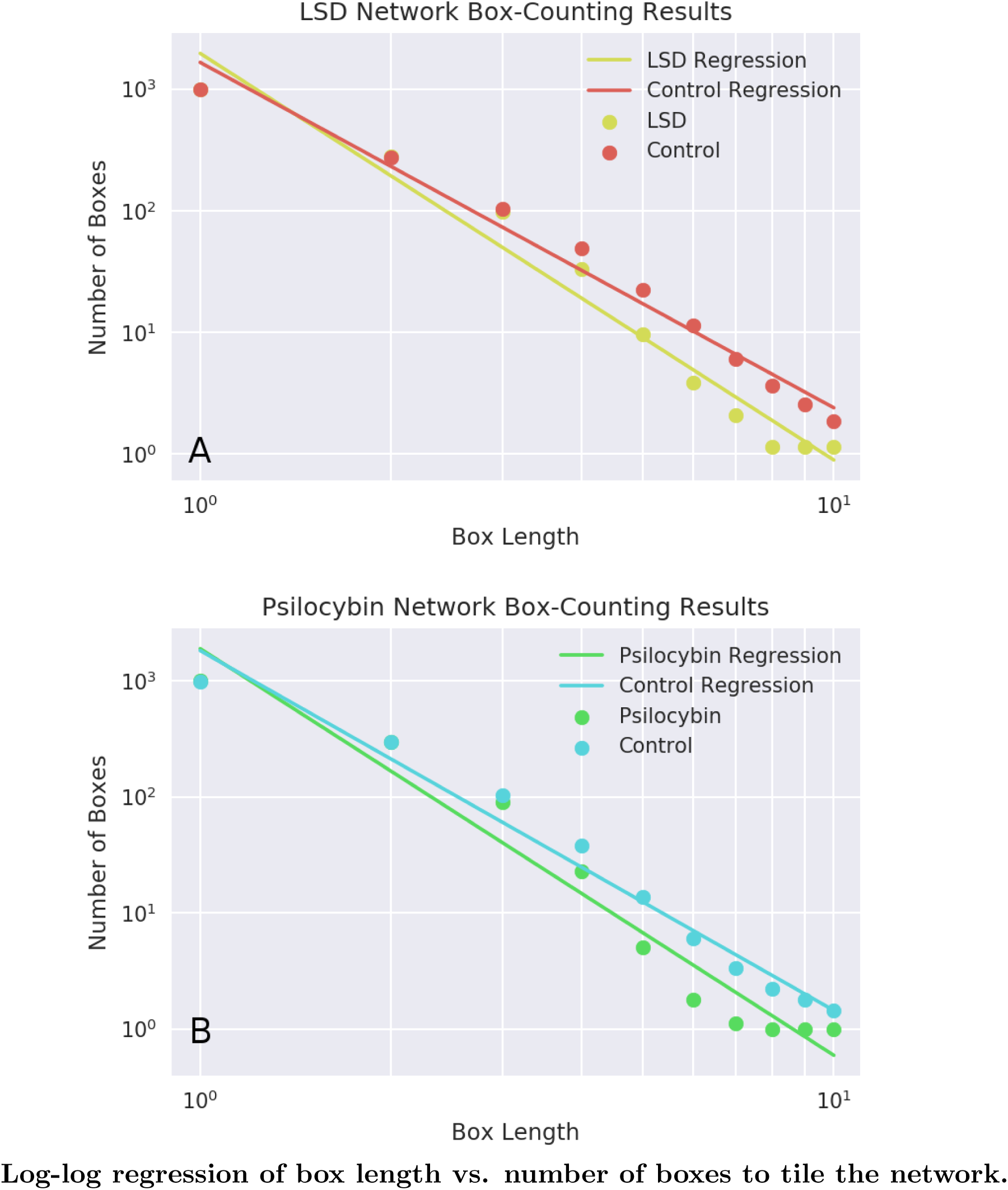
Here is the derivation of the fractal dimension for the LSD and psilocybin tests. For a range of integer-valued box-lengths ({1,2,…,10}), the minimum number of boxes of that length necessary to tile a 1,000-ROI functional connectivity measure is calculated. If the log-transformed values display a linear relationship, that is evidence of a power-law distribution, and the slope characterizes the dimension of the network. Here, each point is th the average number of boxes across all subjects (n=15) in that condition, for each box length. A steeper slope corresponds to a higher fractal dimension, which is associated with a more complex system. Note the log-log axes.

These results are consistent with the EBH, which posits that the properties of criticality will increase during psychedelic states [10]. These results are also consistent with the hypothesis that the changes in brain activity induced by LSD are very similar to the changes induced by psilocybin, which is unsurprising given their shared serotonergic pharmacology and the phenomenological similarities between the associated experiences. The difference in base-line fractal dimension [between LSD and psilocybin] is intriguing: we had expected it to be consistent across both datasets, as normal waking consciousness is presumably similar among volunteers in both datasets. We tentatively hypothesize that it may be a result of differences in data acquisition and processing specifications. It may be, however, that the base-line fractal dimension of BOLD signals is not as consistent between populations as we had assumed, and this may be an interesting future direction of exploration.

### 3.2 LSD & Psilocybin BOLD Time-Series Fractal Dimension

The Wilcoxon signed-rank test, when corrected with the Benjamini-Hochberg procedure with an FDR of 5%, found significant differences between the Higuchi fractal dimension of the LSD time-series and placebo time-series (H(3) p-value=0.001), but not between the pre-infusion and post-infusion psilocybin time-series. The mean network fractal dimension for the LSD-condition time-series was 0.91 ± 0.005 and for the placebo condition it was 0.9 ± 0.006. For the post-infusion psilocybin condition, the mean network fractal dimension of the BOLD time-series was 1.03 ± 0.015, while for the pre-infusion condition it was 1.02 ± 0.009. For visualization of the global Higuchi fractal dimension for the LSD versus control conditions, see Figure 5A, and for visualization of the global Higuchi fractal dimension for the psilocybin versus control conditions, see Figure 5B.

**Figure 5:**
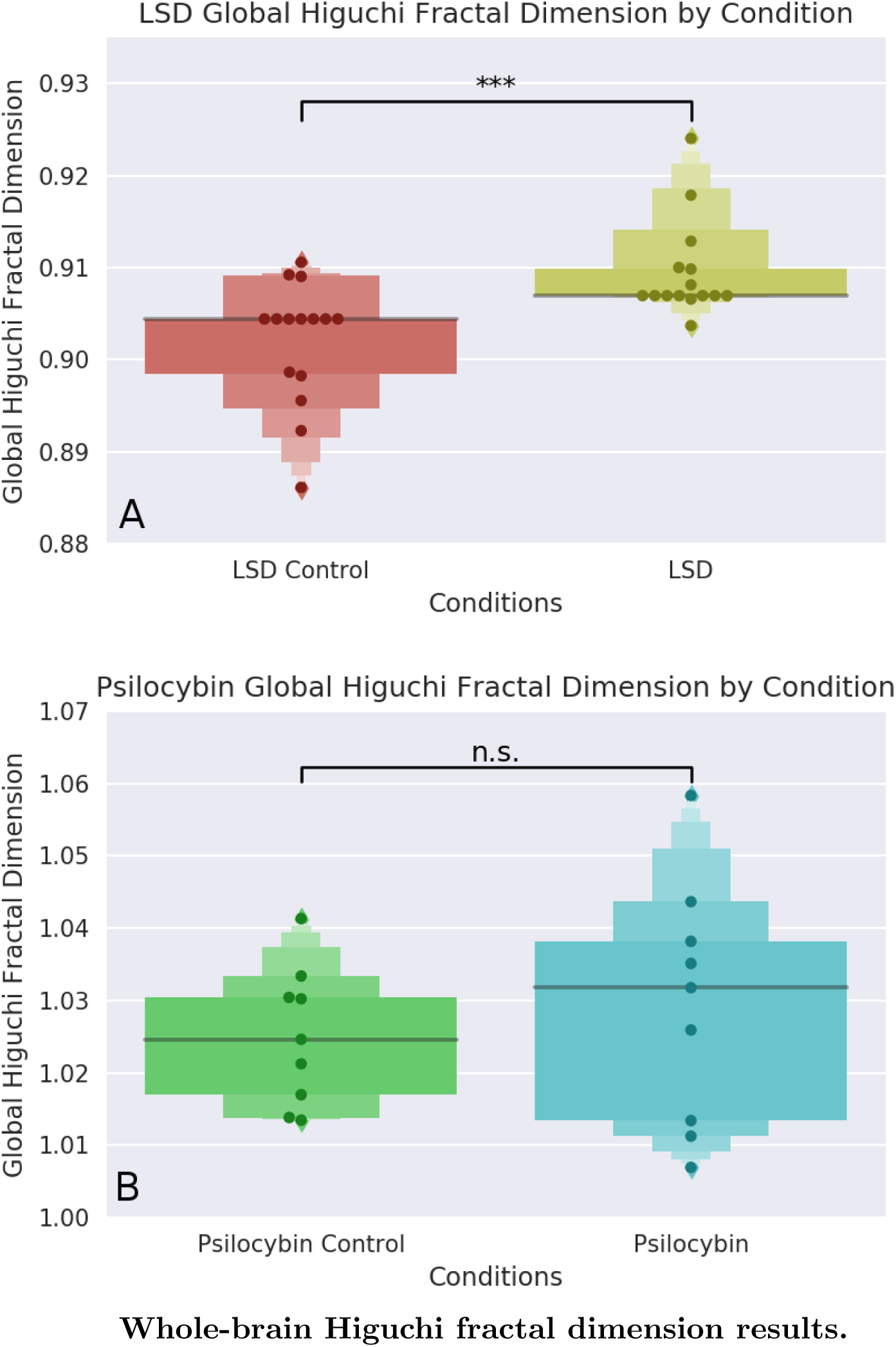
The average Higuchi fractal dimension of BOLD time-series from every one of the 1,000 ROIs used in the Network Fractal Dimension section. Plot A corresponds to the LSD vs. LSD Control condition, Plot B corresponds to the Psilocybin vs. Psilocybin Control condition. For each time-series, the fractal dimension was calculated using a *k*_*max*_ = 64. While the effect size is small in absolute terms, given the small range that the fractal dimension of a time-series usually falls, it remains highly significant.

These results suggest that, at least for the LSD condition, the activity of the brain tends towards increased fractal character in the temporal as well as spatial dimension. This is consistent with the EBH and serves as validation of the network fractal dimension results reported above. The difference between the averages between the two non-drug conditions (placebo condition of the LSD dataset, and the pre-infusion condition of the psilocybin dataset) are most likely explained by the significant difference in the lengths of scans and number of time-points the algorithm was fed. To test this, we re-ran the Higuchi fractal dimension analysis on LSD signals that had been truncated to be the same length as as the psilocybin time-series (100 samples), and found that there was no longer a significant difference between the drug and control conditions. We take this as evidence that the lack of significant difference between psilocybin and control conditions cannot be attributed to the drug directly but rather, may be reflective of a fundamental limitation in the utility of the Higuchi algorithm when working with sparse datasets.

We found a significant correlation between network fractal dimension and temporal fractal di-mension in the LSD condition (*ρ* = 0.49, p-value = 0.006), however, we did not find a significant correlation between the two metrics in the psilocybin conditions (*ρ* = 0.31, p-value = 0.21). For visualization, see Figure 6.

**Figure 6:**
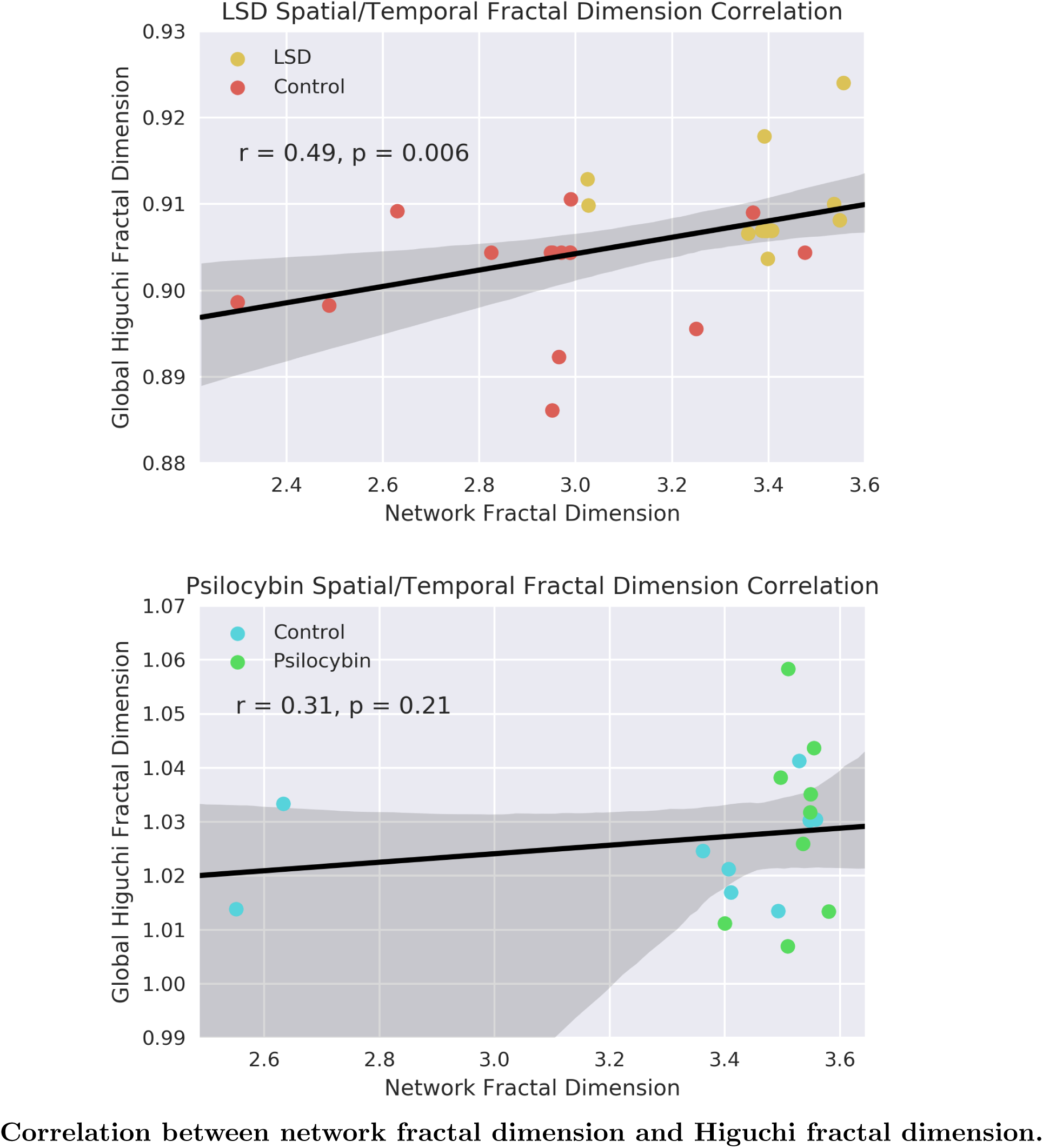
Correlation between network fractal dimension and global higuchi fractal dimension in the LSD and psilocybin conditions. In the LSD dataset, there was a significant, positive correlation between the two measures of fractal dimension (r=0.49, p-value =0.006) that was not apparent in the psilocybin dataset (r=0.23, p-value = n.s.). As previously discussed, we believe this is reflective of the short length of the psilocybin time-series relative to the LSD scans.

#### 3.2.1 Localizing Time-Series Fractal Dimension to Sub-Networks

To take advantage of the fact that the Higuchi method of calculating fractal dimension works on one time-series at a time, we were able to test whether any specific sub-networks of the brain displayed any changes in the fractal-dimension of the associated time-series. For the psilocybin condition, only one significant difference in the fractal dimension of BOLD time-series was found: the fractal dimension increased in the dorsal attentional network, at the edge of significance (H(6), p-value = 0.05). In light of our suspicion that the psilocybin time-series are too short for meaningful Higuchi analysis, we strongly feel that these results should be replicated, using either longer fMRI scans, or, ideally, MEG or EEG data. For a table of the Higuchi fractal dimensions for each network tested in the psilocybin condition, see Table 1.

**Table 1:** Highuchi fractal dimension of BOLD time-series from specific sub-networks in the Psilocybin vs. Control condition

| Sub-Network | Condition | BOLD Fractal Dimension | $p$-Value |
| --- | --- | --- | --- |
| Default-Mode Network | Control | $1.023 \pm 0.016$ | W(14) |
| | Psilocybin | $1.032 \pm 0.017$ | $p = 0.31$ |
| Limbic Network | Control | $1.034 \pm 0.017$ | W(13) |
| | Psilocybin | $1.044 \pm 0.014$ | $p = 0.26$ |
| Fronto-Parietal Network | Control | $1.022 \pm 0.021$ | W(17) |
| | Psilocybin | $1.03 \pm 0.018$ | $p = 0.51$ |
| Somato-Motor Network | Control | $1.031 \pm 0.017$ | W(21) |
| | Psilocybin | $1.028 \pm 0.016$ | $p = 0.86$ |
| Ventral-Attentional Network | Control | $1.031 \pm 0.018$ | W(21) |
| | Psilocybin | $1.033 \pm 0.02$ | $p = 0.86$ |
| Dorsal-Attentional Network * | Control | $1.013 \pm 0.023$ | W(6) |
| | Psilocybin | $1.027 \pm 0.024$ | $p = 0.05$ |
| Visual Network | Control | $1.024 \pm 0.025$ | W(17) |
| | Psilocybin | $1.021 \pm 0.027$ | $p = 0.51$ |

For the LSD condition, compared to the placebo condition, we found significant increases in fractal dimension under LSD in the fronto-parietal network (H(4), p-value = 0.001), in the dorsal-attentional network (H(0), p-value=0.0005), and the visual network (H(4), p-value=0.001). For a table of the Higuchi fractal dimensions for each network tested in the LSD condition, see Table 2.

**Table 2:**
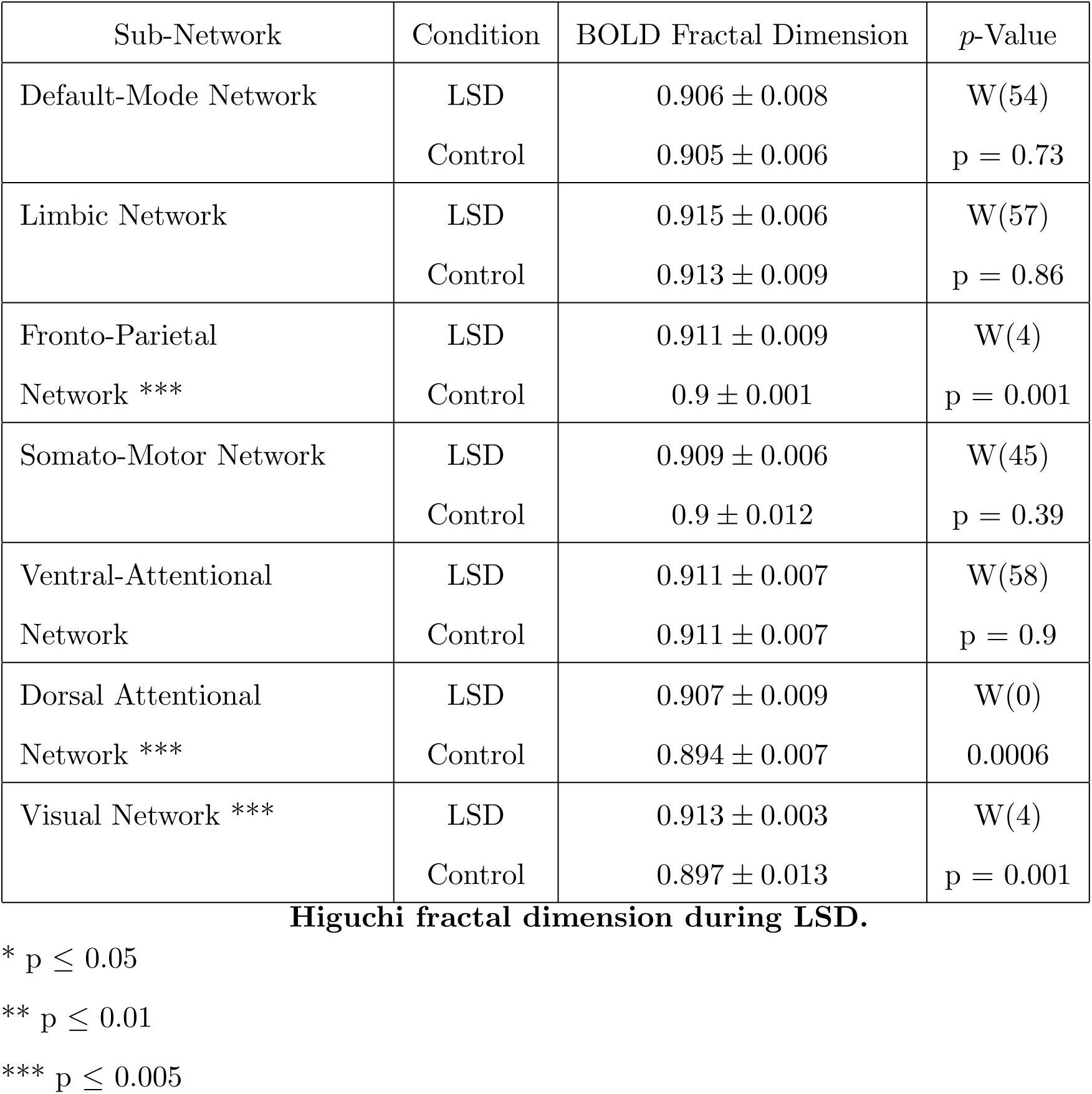
Highuchi fractal dimension of BOLD time-series from specific sub-networks in the LSD vs. Control condition

| Sub-Network | Condition | BOLD Fractal Dimension | *p*-Value |
| --- | --- | --- | --- |
| Default-Mode Network | LSD | $0.906 \pm 0.008$ | W(54) |
| | Control | $0.905 \pm 0.006$ | $p = 0.73$ |
| Limbic Network | LSD | $0.915 \pm 0.006$ | W(57) |
| | Control | $0.913 \pm 0.009$ | $p = 0.86$ |
| Fronto-Parietal<br>Network *** | LSD | $0.911 \pm 0.009$ | W(4) |
| | Control | $0.9 \pm 0.001$ | $p = 0.001$ |
| Somato-Motor Network | LSD | $0.909 \pm 0.006$ | W(45) |
| | Control | $0.9 \pm 0.012$ | $p = 0.39$ |
| Ventral-Attentional<br>Network | LSD | $0.911 \pm 0.007$ | W(58) |
| | Control | $0.911 \pm 0.007$ | $p = 0.9$ |
| Dorsal Attentional<br>Network *** | LSD | $0.907 \pm 0.009$ | W(0) |
| | Control | $0.894 \pm 0.007$ | 0.0006 |
| Visual Network *** | LSD | $0.913 \pm 0.003$ | W(4) |
| | Control | $0.897 \pm 0.013$ | $p = 0.001$ |

The significant increase in the dorsal-attentional network in both the LSD and psilocybin conditions suggests that this finding may be more robust than the increases in the fronto-parietal network or visual network that appear to be unique to LSD. An increase in the complexity of activity in the visual system under LSD is somewhat unsurprising, although why this did not appear in psilocybin is unclear (under the psilocybin condition the mean complexity in the visual system did increase relative to the pre-infusion condition, although this was not significant).

## 4 Discussion

Here, we report that, using a Compact-Box Burning algorithm [49], the fractal dimension of high-resolution cortical functional connectivity networks is increased under the influence of both psilocybin and LSD, both serotonergic psychedelic compounds, and that the fractal dimension of the BOLD time-series is increased by LSD, but not psilocybin. Furthermore, for both LSD and psilocybin, we were able to show a significant increase in the fractal dimension of the BOLD time-series in the brain regions generally thought to make up the dorsal-attentional network. These results suggest that psychedelic drugs increase the fractal character of brain activity in both temporal (as measured by Higuchi fractal dimension), and spatial domains (as measured by the Compact-Box burning algorithm). We interpret this result as an indicator that, under the influence of psychedelics, the brain moves towards a region of criticality [26, 27, 28], as fractal qualities emerge as the system nears a tipping point, or transition zone, from one phase into another [67]. This is in keeping with the predictions of the Entropic Brain Hypothesis (EBH), which hypothesizes that the level and quality of consciousness changes as the brain evolves towards the zone of criticality, between distinct phases [10, 4]. Our results also line up nicely with other attempts to quantify the complexity of brain activity under psychedelics, which have generally reported increases in entropy relative to an unaltered baseline [17, 16, 68, 20, 18].

One question that remains unanswered is what exactly the qualitative differences between those two phases might be: as was previously mentioned, the EBH intuitively lends itself to an Ising-like interpretation, where the critical moment partitions a low-entropy state and a more random, high-entropy phase, although this raises difficult questions about how that phase may present in a living, biological system. The critical Ising model has been used as a model for brain activity and may capture instrinic properties of neural self-organization [29, 30, 31] An alternative model of criticality may be one of a branching process [69], where in the sub-critical regime the propagation of a branch is guaranteed to halt eventually, while in the super-critical regime, the branch flourishes, and at the point of criticality, the process branches into fractal patterns [70]. Simulations of neural networks suggest that super-critical behaviour should be epileptiform in nature [71], but psychedelics, on their own, do not typically induce seizures [72] (although collected anecdotal reports have suggested that LSD, in combination with lithium can increase the risk of seizures [73], as can the the psilocybin analogue 5-methoxy-dimethyltryptamine [74]).

One interesting direction of research these results suggest is an analysis of whether the fractal dimension of a network, such as those explored here, encodes any information about the ability of that network to integrate information, a key issue of Integrated Information theory (IIT), [12, 13, 14]. Simulations of small networks have found that the topology of a network can have implications for its capacity to act as an integrator of information [75]. In the cited simulation, network complexity was highest in a modular network based on the architecture of the visual system compared to a simpler, less integrated, network of the same type, or a network with a random distribution of connections. This idea of balancing integration and modularity recalls the findings by Gallos et al., that the fractal quality of functional connectivity networks plays a role in balancing these two competing topologies in a manner optimal for computation [44]. While it is computationally infeasible to do a crude calculation of integrated information for any non-trivial neural system due to the explosive growth in the number of computations involved, methods of estimating the value have been developed [76, 77], and so, using the fractal dimension analysis method described here, it should be possible to begin to explore whether there is a relationship between the fractal dimension of a system and it’s ability to integrate information. Recently it has been shown that, in models of self-organizing, critical systems, such as Abelian sandpiles (which naturally tend towards critical states due to repeated build-up and relaxation of energy as the system evolves), critical behavior was surprisingly good at optimizing certain hard computational problems on graphs [78], suggesting that criticality may underlie some of the brain’s own computational abilities.

While the theoretical implications for these results in the context of the EBH are interesting on their own, we also try to ground these results in the current literature concerning the neurobiology of psychedelic drugs. All serotonergic psychedelics (eg: LSD, mescaline, psilocybin) share agonist activity at the 5-HT2A receptor [79], a metabotropic serotonin receptor known to be involved in modulating a variety of behaviours. While the 5-HT2Ar is widely expressed in the CNS, a specific population localized to Layer V pyramidal cells in the neocortex is both necessary and sufficient to induce psychedelic effects [80]. These Layer V pyramidal neurons serve as ‘outputs’ from one region of the cortex to another [81], and the 5-HT2Ar acts as an excitatory receptor, decreasing polarization and increasing the probability that a given neuron will fire [82, 83]. This suggests a primitive model of 5-HT2Ar’s role in neural information processing: on Layer V pyramidal neurons, the 5-HT2Ar serves as a kind of ‘information gate’. When a psychedelic is introduced to the brain, it binds to the 5-HT2Ar, exciting the associated pyramidal neuron and decreasing the threshold required to successfully transmit information through the neuron. During normal waking consciousness, areas of the brain that are physically connected by Layer V pyramidal neurons may not be functionally connected because the signal threshold required to trigger an action potential is too high but when a psychedelic is introduced, that threshold goes down allowing novel patterns of information flow to occur. This perspective also recalls the branching process discussed above [69]. In this case, increasing the probability of a pyramidal neuron firing may be analogous to increasing the branching ratio *σ*, which, if *σ* is normally sub-critical, would bring the process closer to the critical value of *σ*_*c*_. As networks with fractal topology are related to the trees generated by critical branching processes [70], this may be a fruitful area to explore further.

It is difficult to interpret the increase in the fractal dimension of the BOLD time-series in the dorsal-attentional network. This network is generally thought to be involved in a variety of processes related to visual processing of the environment, such as attending to the orientation of objects in space, visual feature-based attention, and biasing visual perception in response to cues [84]. It was originally proposed to be involved with top-down, conscious allocation of attention to environmental objects [85]. Human studies with psilocybin have found that exposure to the psychedelic reduces attentional tracking ability, and the proposed mechanism given was that psilocybin reduced the ability of the brain to filter out irrelevant or distracting stimuli [86]. This is consistent with findings that psychedelics attenuate sensory-gating functions in a manner reminiscent of patients with schizophrenia [87, 88].

The finding that LSD increased the fractal dimension of BOLD signals in the fronto-parietal network is consistent with previous findings that global increases in the functional connectivity density induced by LSD overlap with brain regions commonly assigned to the FP network [89]. We did not, however find significant changes in the complexity of signals from nodes commonly assigned to the Default Mode Network (DMN), which ran counter to our initial hypothesis. Many neuroimaging studies of psilocybin and LSD have found associations between changes in DMN activity and the phenomonology of the psychedelic experience [59, 58, 89, 90]. We hypothesize that this discrepancy might be explained by the sheer number of nodes assigned to the DMN (212 nodes in total): because the signal from every node was weighted equally, it is possible that peripheral nodes assigned to the DMN by our parcellation may not have been significantly effected, thus obscuring a real effect only present in a subset of DMN nodes. Validation with a smaller atlas or more conservative assignment of nodes may yet find an effect in the DMN (although a smaller atlas would preclude the NFD analysis).

Finally, the increased complexity of BOLD signals in the visual network under LSD is interesting, although perhaps unsurprising given the fantastically visual nature of the psychedelic experience. It has already been established that LSD alters functional connectivity of visual cortices in humans [91], and EEG analysis of LSD users post-experience has found alterations to the coherence of signals in visual areas thought to be associated with the experience of hallucinations [92]. It has been suggested that the qualitative nature of psychedelic imagery may be informative about the structure and layout of the visual system [93], and so we propose that this may be a particularly fruitful avenue of psychedelic research going forward.

This study has several limitations that are worth considering. The first is the comparatively small size of the psilocybin sample (n=9), which means that it is harder to trust the replicability of the present findings than if the sample had been larger. Second, the Higuchi fractal dimension is not frequently used on BOLD signals, as the number of samples in each time-series is far lower than it is for EEG or MEG, resulting in a less robust analysis. In the case of psilocybin, the time-series may be so too short too produce Higuchi fractal dimension values of any reliability. In light of this, replication with EEG or MEG data should be a priority before these results are considered strong. Simultaneous EEG-fMRI recordings under a psychedelic would be particularly informative as it would enable us to test the relationship between fractal dimension recorded across modalities. Third, the parcellation resolution used here (1000 ROIs), which is considerably larger than many commonly-used parcellations is still smaller than would be desired for a truly comprehensive analysis of fractal dimension of functional connectivity networks, and so future analysis with a higher resolution cortical parcellation is needed. Future studies comparing different psychedelics, like LSD and psilocybin, should also strive to ensure some kind of dose-equivalence: given the nature of the datasets, it was not possible to ensure that the subjective intensities of the LSD and psilocybin experiences volunteers underwent was equivalent, and this may be reflected in the differences in results. To control for this, it would be valuable to have a universal, standardized measure of subjective experience such as the ASC questionnaire [94], with graded doses for a variety of drugs, such as psilocybin, LSD, mescaline, etc. This would allow researchers the ability to more fully explore the commonalities, and differences between individual psychedelic compounds.

Finally, it is unclear what the functional, psychological implications of increased fractal properties of brain activity and network organization are. Particularly profound subjective experiences under moderate-high doses of psychedelics are a highly reliable observation. Although there are clear differences in the specific vocabulary and intellectual framing used to describe and depict these experiences, variously referred to as peak experiences by some [95] and mystical-type experiences by others [96], there is clear consensus that the phenomenology of the experience itself is fundamental, and that its nature is often felt as exceptional in terms of both novelty and perceived meaning [7]. Based on the present studys findings it is reasonable to speculate that the changes observed here, which are consistent with a system nearing criticality, may relate in some way to these profound subjective effects of psychedelics which include: exceptional sensitivity to environmental perturbation [97] and a sense of oneness or connectedness [98] including a sense of attunement to or aligned with nature [99, 97], referred to as the unitive experience [100] and thought to be a principal component of the peak/mystical-type experience [101]. The original EBH speculated that a closer tuning of brain activity to criticality may better reflect the ubiquitous criticality evident throughout the natural world and thus account for the subjective feeling of being better attuned to nature [10]. Future work is now required to assess these speculative ideas and test the nature of the associations with greater specificity. This will demand improvements in sampling of the subjective experience [102] as much, if not more so, than improvements in the sampling of brain activity. Improving our understanding of the brain basis of the psychedelic experience may have implications for our understanding of how these compounds might be best utilized, e.g. as aides to psychological development and therapy [10] as well as how they may model specific aspects of psychosis [103, 104].

## 5 Conclusions

In this study we report that, under the influence of two serotonergic psychedelics: LSD and psilocybin, the fractal dimension of cortical functional connectivity networks is significantly increased. Under LSD, the fractal dimension of BOLD time-series is also significantly increased, while psilocybin shows a non-significant increase as well. These results are in line with previously published research suggesting that psychedelics increase the complexity of brain activity, and the specific measures used here may be a particularly useful tool for understanding how consciousness changes as the brain approaches criticality. We were able to show that, under both LSD and psilocybin, the fractal dimension of BOLD time-series from regions assigned to the dorsal-attentional network was increased. These findings show that psychedelics increase the fractal dimension of brain activity in both spatial and temporal domains and have implications for the study of consciousness and the neurobiology the psychedelic experience.

## Supporting information

Supplemental data 1, LSD fractal dimensions

Supplemental data 2, psilocybin fractal dimensions

## Conflict of Interest Statement

The authors report no personal or financial conflicts of interest related to the research reported herein.

### Acknowledgements

The authors would like to thank the Cambridge University Department of Anaesthetics. We would like to specifically thank: Ioannis Pappas, Michael Craig, Dian Lu, and Andrea Luppi for their support. We would also like to thank Dr. Fernando Rosas for detailed feedback and insights. DK Menon is funded by the MRC, the NIHR Cambridge Biomedical Centre, and an NIHR Senior Investigator Award, and EA Stamatakis is funded by the Stephen Erskine Fellowship Queens College Cambridge. L Roseman has been supported by an Imperial President’s Scholarship. Robin Carhart-Harris has been supported by the Beckley Foundation and is now supported by the Alex Mosley Charitable Trust and Ad Astria Chandaria Foundation. The original LSD research was completed with the support of a Walacea.com crowd-funding campaign and the Beckley Foundation. The psilocybin research was completed with the support of the Beckley Foundation and with additional support from the Neuro-psychoanalysis Foundation, Multidisplinary Association for Psychedelic Studies, and the Heffter Research Institute.

## Author Contributions

TFV, DKM and EAS carried out the research reported here. RC-H and LR designed the initial experiments, collected and preprocessed the data. TFV designed and performed the fractal analyses with feedback from EAS. TFV wrote the paper with feedback from all co-authors.

